# Suppressed vascular Rho-kinase activation is a protective cardiovascular mechanism in obese female mice

**DOI:** 10.1101/2023.04.12.536489

**Authors:** Gabriela Souza Barbosa, Rafael M Costa, Wanessa MC Awata, Shubhnita Singh, Juliano V Alves, Ariane Bruder-Nascimento, Camila Renata Corrêa, Thiago Bruder-Nascimento

## Abstract

**Background:** Obesity is the number one cardiovascular risk factor for both men and women and is a complex condition. Although a sex dimorphism on vascular function has already been noted, the underlying processes remain unclear. The Rho-kinase pathway has a unique role in controlling vascular tone, and in obese male mice, hyperactivation of this system results in worsened vascular constriction. We investigated whether female mice exhibit decreased Rho-kinase activation as a protective mechanism in obesity.

**Methods:** We exposed male and female mice to a high-fat diet (HFD) for 12 weeks. At the end, energy expenditure, glucose tolerance, adipose tissue inflammation, and vascular function were investigated.

**Results:** Male mice were more sensitive to HFD-induced body weight gain, glucose tolerance, and inflammation than female mice. After establishing obesity, female mice demonstrated increase in energy expenditure, characterized by an increase in heat, whereas male mice did not. Interestingly, obese female mice, but not male, displayed attenuated vascular contractility to different agonists, such difference was blunted by inhibition of Rho-kinase. Finally, aortae from obese female mice, but not male, responded prematurely to Rho-kinase inhibitor, which was accompanied by a suppressed Rho-kinase activation, measured by western blot.

**Conclusion:** In obesity, female mice demonstrate a vascular protective mechanism – suppression of vascular Rho-kinase – to minimize the cardiovascular risk associated with obesity, whereas male mice do not generate any adaptive response. Future investigations can help to understand how Rho-kinase becomes suppressed in female during obesity.

## Introduction

Obesity is a multifactorial disease with a complex pathogenesis associated with psychosocial socioeconomic, biological, and environmental mechanisms^1^. Obesity is a major public health issue that has been on the rise in the world. The World Health Organization (WHO) reports that the global obesity rate has nearly tripled since 1975, and the obesity epidemic is now well-established^2^. Obesity increases the risk of a number of metabolic abnormalities, including type 2 diabetes, hypertension, inflammation, and dyslipidemia, which are major risk factors of vascular injury and cardiovascular disease (CVD)^1-3^. Therefore, to enhance the quality of life and lower the mortality linked to this condition, it is essential to understand the vascular pathways causing cardiovascular disease in obesity.

Obesity is a leading risk factor for CVD and a major health burden in male and female^4-6^. However, the sex-discrepant mechanism is implicated in obesity-associated CVD^7-9^. Sex hormones, as well as sex chromosomes themselves can cooperate to the development of obesity, glucose metabolism, and vascular function regulation^5-11^. In 2017^5^, we showed that females display a slower body weight gain compared to male mice under high fat diet (HFD), which is followed by protection against obesity-induced sympathetic activation and changes in adrenergic vascular contractility. Although we observed a difference in vascular response in male and female mice, we did not identify the underlying vascular mechanisms.

Rho-kinase, a downstream target in the RhoA-linked pathway, is formerly identified as an effector of the small GTPase Rho. In the vasculature, RhoA-linked pathway can determine motility, morphology, polarity, cell division, gene expression, and cellular contraction. Rho-kinase promotes vascular contraction via a complex and extensive network between RhoA, Rho-kinase or ROCK (ROCKα/ROCK2), ROCKβ/ROCK1, myosin phosphatase target subunit 1 (MYPT1), and myosin light chain (MLC)^12-14^. Increased vascular tone in obesity has been previously demonstrated to be Rho-kinase overactivation-dependent in males^15, 16^, but whether females present the same Rho-kinase overactivation dependence on vascular contraction in obesity is still to be determined.

In addition to characterizing the body weight gain, glucose sensitivity, and energy expenditure in male and female mice under HFD treatment, this study also sought to understand the difference in vascular contractility between male and female mice in obesity. Therefore, we tested the hypothesis that female mice display a suppressed Rho-kinase activation as a compensatory mechanism to attenuate vascular contractility in obesity.

## Methods

### Mice

Male and female C57Bl/6 mice (6-8 weeks of age) were divided into 4 groups and fed either a normal diet (ND; Research Diets, D12328, Carbohydrate 73%, Fat 11%, and Protein 16% kcal) or a high-fat diet (HFD; Research Diets, D12492, 60% of fat calories; 20% of protein and 20% of carbohydrate) ad libitum. Tap water was provided ad libitum. Mice were monitored for 12 weeks. Body weight was measured weekly. Mice were housed in an American Association of Laboratory Animal Care–approved animal care facility in the Rangos Research Building at the Children’s Hospital of Pittsburgh of the University of Pittsburgh. Institutional Animal Care and Use Committee approved all protocols (IACUC protocols # 19065333 and 22061179). All experiments were performed in accordance with Guide Laboratory Animals for The Care and Use of Laboratory Animals. At the end of the experiments gonadal, retroperitoneal, visceral, subcutaneous, subscapular brown adipose tissue, heart, liver, and kidneys were isolated and weighed for adiposity and cardio-renal characterization.

### Energy expenditure

The Oxymax Lab Animal Monitoring System (CLAMS, Columbus Instruments, Columbus, OH) was used to determine Heat and Respiratory Exchange Ratio (RER) calculated from CO_2_ production and O_2_ uptake ratio as described before^17^. Mice were placed on CLAMS for 2 days of acclimatization, then the parameters mentioned above were recorded for 72 hours. Area under curve from 72h record was used to determine any difference.

### Intraperitoneal glucose tolerance test (ipGTT)

IpGTT was performed to evaluate glucose tolerance. Mice were deprived of food for 12h. Blood sample was collected from the caudal vein immediately before (baseline, t_0_) and after (t_15_, t_30_, t_60_, t_90_, t_120_ min) administration of 2 g of glucose/kg by intraperitoneal injection. Glucose levels were determined using a glucose analyzer (Accu-Check, Roche Diagnostics) as previously described^18^.

### Vascular remodeling

Mice were euthanized for aortae harvest and perfused with cold phosphate-buffered saline (PBS). Aortae were collected and placed in a 4% paraformaldehyde (PFA) solution for histology analysis. After 12h in PFA, tissues were placed in 70% ethanol until the day of preparing the samples for histology. Aortae were embedded in paraffin, then samples were sectioned and stained with hematoxylin and eosin (H&E) to analyze the vascular remodeling and structure.

### Adipose tissue inflammation

mRNA from gonadal fat was extracted using RNeasy Mini Kit (Quiagen, Germantown, MD – USA). Complementary DNA (cDNA) was generated by reverse transcription polymerase chain reaction (RT-PCR) with SuperScript III (Thermo Fisher Waltham, MA USA). Reverse transcription was performed at 58 °C for 50 min; the enzyme was heat inactivated at 85 °C for 5 min, and real-time quantitative RT-PCR was performed with the PowerTrackTM SYBR Green Master Mix (Thermo Fisher, Waltham, MA USA). Sequences of genes as listed in supplementary table 1. Experiments were performed in a QuantStudioTM 5 Real-Time PCR System, 384-well (Thermo Fisher, Waltham, MA USA). Data were quantified by 2ΔΔ Ct and are presented by fold changes indicative of either upregulation or downregulation.

### Western blot

Aortic protein was extracted using radioimmunoprecipitation assay buffer (RIPA) buffer (30mM HEPES, pH 7.4,150mM NaCl, 1% Nonidet P-40, 0.5% sodium deoxycholate, 0.1% sodium dodecyl sulfate, 5mM EDTA, 1mM NaV04, 50mM NaF, 1mM PMSF, 10% pepstatin A, 10 μg/ml leupeptin, and 10 μg/ml aprotinin). Protein samples were suspended in Laemmli Sample Buffer supplemented with 2-Mercaptoethanol (β-mercaptoethanol) (BioRad Hercules, California – USA). Then, proteins were separated by electrophoresis on a polyacrylamide gradient gel (BioRad Hercules, California – USA), and transferred to Immobilon-P poly (vinylidene fluoride) membranes. Non-specific binding sites were blocked with 5% skim milk or 1% bovine serum albumin (BSA) in tris-buffered saline solution with tween for 1h at 24 °C. Membranes were then incubated with specific antibodies overnight at 4 °C as described in supplementary table 2. After incubation with secondary antibodies, the enhanced chemiluminescence luminol reagent (SuperSignalTM West Femto Maximum Sensitivity Substrate, Thermo Fisher Waltham, MA, USA) was used for antibody detection.

### Vascular reactivity

Endothelium intact aortic rings were mounted in a wire myograph (Danysh MyoTechnology) for isometric tension recordings with PowerLab software (AD Instruments) as described^18-21^. Briefly, rings (2mm) were placed in tissue baths containing warmed (37 °C), aerated (95% O2, 5% CO2) Krebs Henseleit Solution: (in mM: 130 NaCl, 4.7 KCl, 1.17 MgSO4, 0.03 EDTA, 1.6 CaCl 2, 14.9 NaHCO3, 1.18 KH2PO4, and 5.5 glucose) and after 30 min of stabilization, arteries were incubated with KCl (60mM) to test the sample viability. Then, the following concentration response curves (CRC) were performed: Phenylephrine and thromboxane analogue (U46619). To study the role of Rho-Kinase pathway, we inhibited ROCK with Y-27632 (100uM) and performed CRC to U46619 or built CRC (0.01nM to 100uM) of relaxation to Y-27632 on top of a pre-constriction evoked by U46619 (10nM).

### Statistical analysis

Our aim was to determine the impact of HFD on male and female mice, thus we used Student’s t-test to determine any difference between ND and HFD in both sexes. The vascular contractility data are expressed in millinewton (mN). The concentration-response curves were fitted by nonlinear regression analysis. Maximal response (Emax) was determined. Analyses were performed using Prism 9.0 software (GraphPad). A difference was considered statistically significant when P ≤ 0.05.

## Results

### Female mice present an attenuated body weight gain, energy expenditure impairment, and glucose tolerance in HFD-induced obesity model

Firstly, we investigated if HFD promotes obesity and impairs energy expenditure and glucose sensitive in male and female mice. By measuring body weight gain and fresh weight of different adipose tissue depots (Fig.1A and C and table 1), we observed that HFD induced obesity in male and female, however female mice were more resistance to obesity appearance. Such an increase in adiposity was followed by a significant enhancement in Ki67 (a marker of proliferation) in gonadal fat in males, but not in females (Fig.1B and 1D).

**Table 1.** Characterization of adiposity and cardiorenal system of male and female mice exposed to normal diet (ND) or high-fat diet (HFD)

| Variable | Groups |  |  |  |
| --- | --- | --- | --- | --- |
|  | Male ND | Male HFD | Female ND | Female HFD |
| Initial body mass (g) | 24.9±0.9 | 26.1±0.42 | 22.9±0.3 | 19.2±0.4 |
| Final body mass (g) | 30.9±1.36 | 48.2±0.9* | 22.8±0.3 | 30.7±2.0<sup>#</sup> |
| Weight gain (g) | 6.0±0.71 | 22.1±0.8* | 1.9±0.3 | 11.5±1.7<sup>#</sup> |
| Gonadal Adipose Tissue (g) | 0.25±0.02 | 1.07±0.09* | 0.20±0.03 | 0.86±0.19<sup>#</sup> |
| Retroperitoneal Adipose Tissue (g) | 0.08±0.01 | 1.09±0.04* | 0.07±0.01 | 0.59±0.12<sup>#</sup> |
| Visceral Adipose Tissue (g) | 0.12±0.03 | 1.01±0.11* | 0.11±0.01 | 0.33±0.07<sup>#</sup> |
| Subcutaneous Adipose Tissue (g) | 0.30±0.05 | 2.09±0.07* | 0.26±0.02* | 1.01±0.16<sup>#</sup> |
| Subscapular Brown Adipose Tissue (g) | 0.07±0.01 | 0.23±0.01* | 0.10±0.01 | 0.12±0.01 |
| Adiposity Index (%) | 2.29±0.26 | 11.23±0.73* | 2.87±0.20 | 8.63±1.16<sup>#</sup> |
| Heart (g) | 0.08±0.01 | 0.10±0.01* | 0.07±0.01 | 0.10±0.02<sup>#</sup> |
| Liver (g) | 0.90±0.04 | 1.80±0.06* | 0.65±0.02 | 0.81±0.05<sup>#</sup> |
| Kidney (g) | 0.24±0.01 | 0.30±0.02* | 0.17±0.01 | 0.21±0.01<sup>#</sup> |

Data are presented as Mean ± SEM. N=4-8. \*P<0.05 vs male ND; <sup>#</sup>P<0.05 vs female ND. Statistic analyzed was performed by comparing ND and HFD within the same sex.

**Figure 1.**
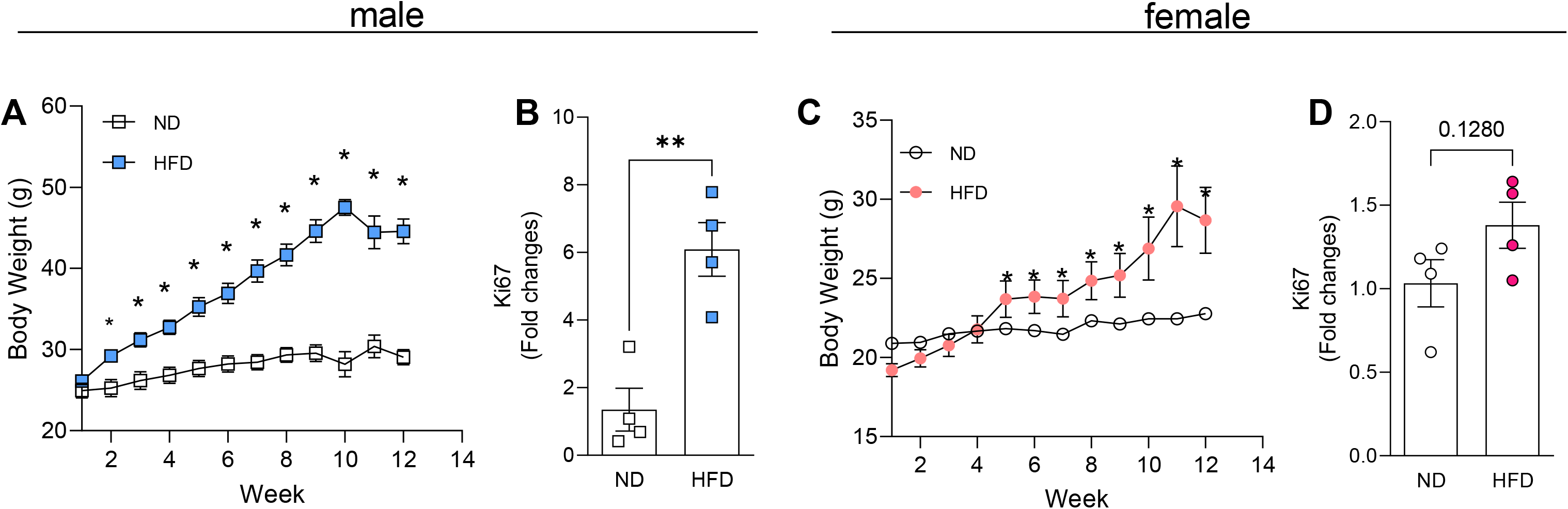
Body weight gain in male and female mice under high fat diet (HFD) treatment. Body weight gain (A and C) and gene proliferation marker in gonadal fat (Ki67, B and D) from male and female mice exposed to HFD for 12. Body weight was analyzed weekly. Ki67 expression was analyzed by RT-PCR. Data are presented as mean ± Standard Error of the Mean (SEM). N=4 for RT-PCR and 8 for body weight gain. *P<0.05 vs normal diet (ND).

Furthermore, we observed that male mice do not present increase in heat after HFD treatment (Fig.2A and B), different from female mice, which demonstrated elevated heat post HFD treatment (Fig. 2E and F). Finally, HFD exposure decreased RER in male and female mice (Fig. 2C-D and 2G-H).

**Figure 2.**
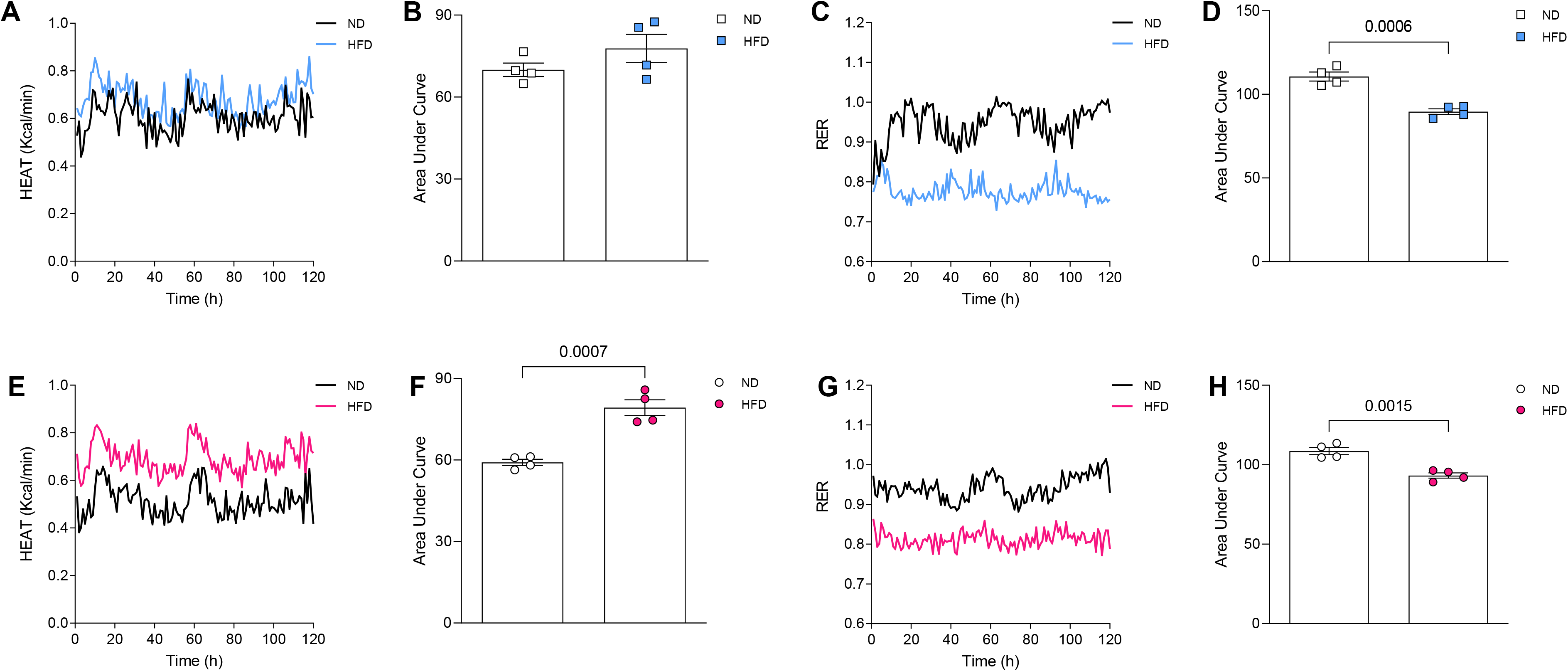
Energy expenditure in male and female mice exposed to high fat diet (HFD). Heat (A, B and E, F) and Respiratory Exchange Ratio (RER) (C, D and G, H) from male and female mice exposed to normal diet (ND) or HFD for 11 weeks. Area under curve data are presented as mean ± Standard Error of the Mean (SEM). N=4. *P<0.05 vs normal diet (ND).

Since obesity is associated with glucose tolerance, we investigated how is the glucose sensitive in male and female mice under HFD treatment, we found that HFD induced glucose tolerance in male and female mice, but male mice appeared to be more resistant to HFD-induced glucose tolerance. (Fig3A-D).

**Figure 3.**
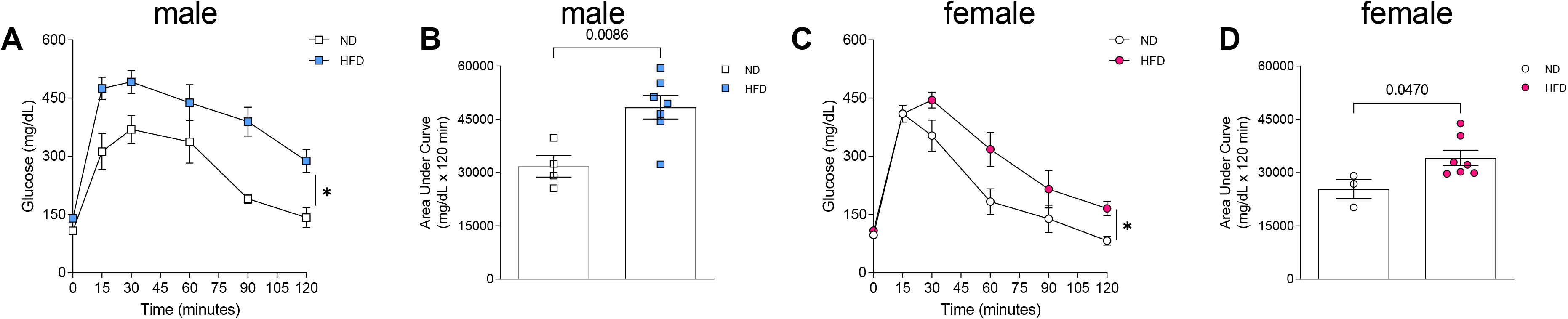
Glucose tolerance in male and female mice exposed to high fat diet (HFD). Intraperitoneal glucose tolerance test (ipGTT) in male (A and B) and female (C and D) mice exposed to normal diet (ND) or HFD for 11 weeks. Data are presented as mean ± Standard Error of the Mean (SEM). N=4. *P<0.05 vs normal diet (ND).

### Female mice are resistant to HFD-induced obesity-associated adipose tissue inflammation

Low-grade inflammation of adipose tissue is a key characteristic of obesity. We investigated, via RT-PCR, the inflammation level in gonadal fat from male and female mice and found that CCR5, ICAM1, VCAM1, and F4/80 (macrophage marker) are elevated only in gonadal fat from males exposed to HFD, whereas TNFα was elevated only in female treated with HFD. Finally, IL6 gene expression was surprisingly decreased in gonadal fat from males treated with HFD (Fig. 4A-F).

**Figure 4.**
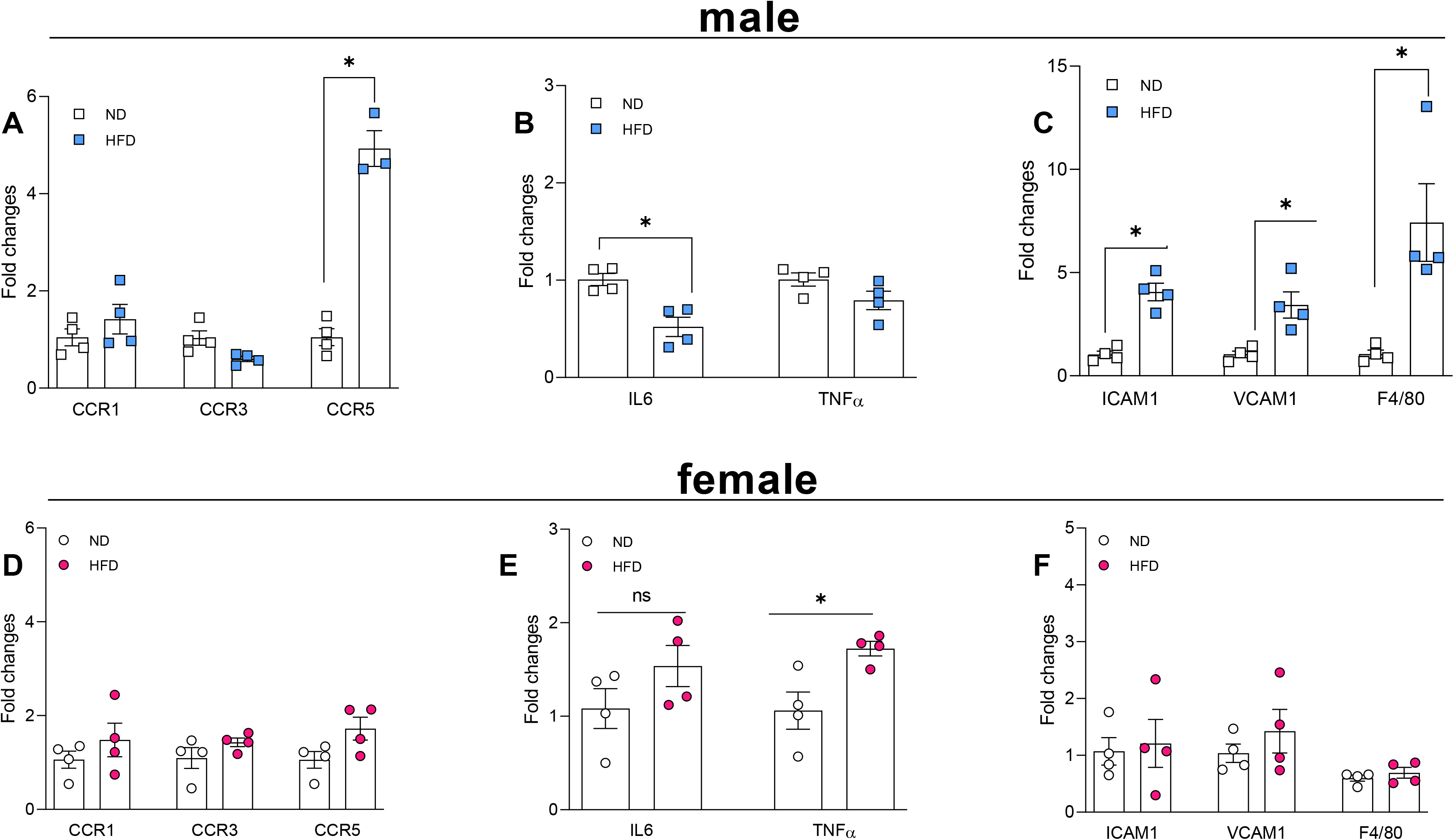
Adipose tissue inflammation in male and female mice exposed to high fat diet (HFD). Chemokines expression (A and D), cytokines (B and E), and adhesion gene and macrophage marker (F4/80) expression (C and F) in gonadal fat from male and female mice exposed to normal diet (ND) or HFD for 12 weeks. Gene expression was analyzed by RT-PCR. Data are presented as mean ± Standard Error of the Mean (SEM). N=4. *P<0.05 vs normal diet (ND).

### Obese female mice demonstrate attenuated vascular contractility with no changes in vascular hypertrophy or contractile protein

Interestingly HFD treatment did not affect the vascular contractility in male mice analyzed by KCl, thromboxane analogue, and phenylephrine responses (Fig. 5A-C), however female mice demonstrated an attenuated vascular contraction to KCl, thromboxane analogue, and phenylephrine (Fig. 5F-H). Finally, changes in vascular response were not dependent on structural modifications or contractile protein content, measured by H&E staining and αSMA amount, respectively (Fig. 5D and E and Fig.5I and J).

**Figure 5.**
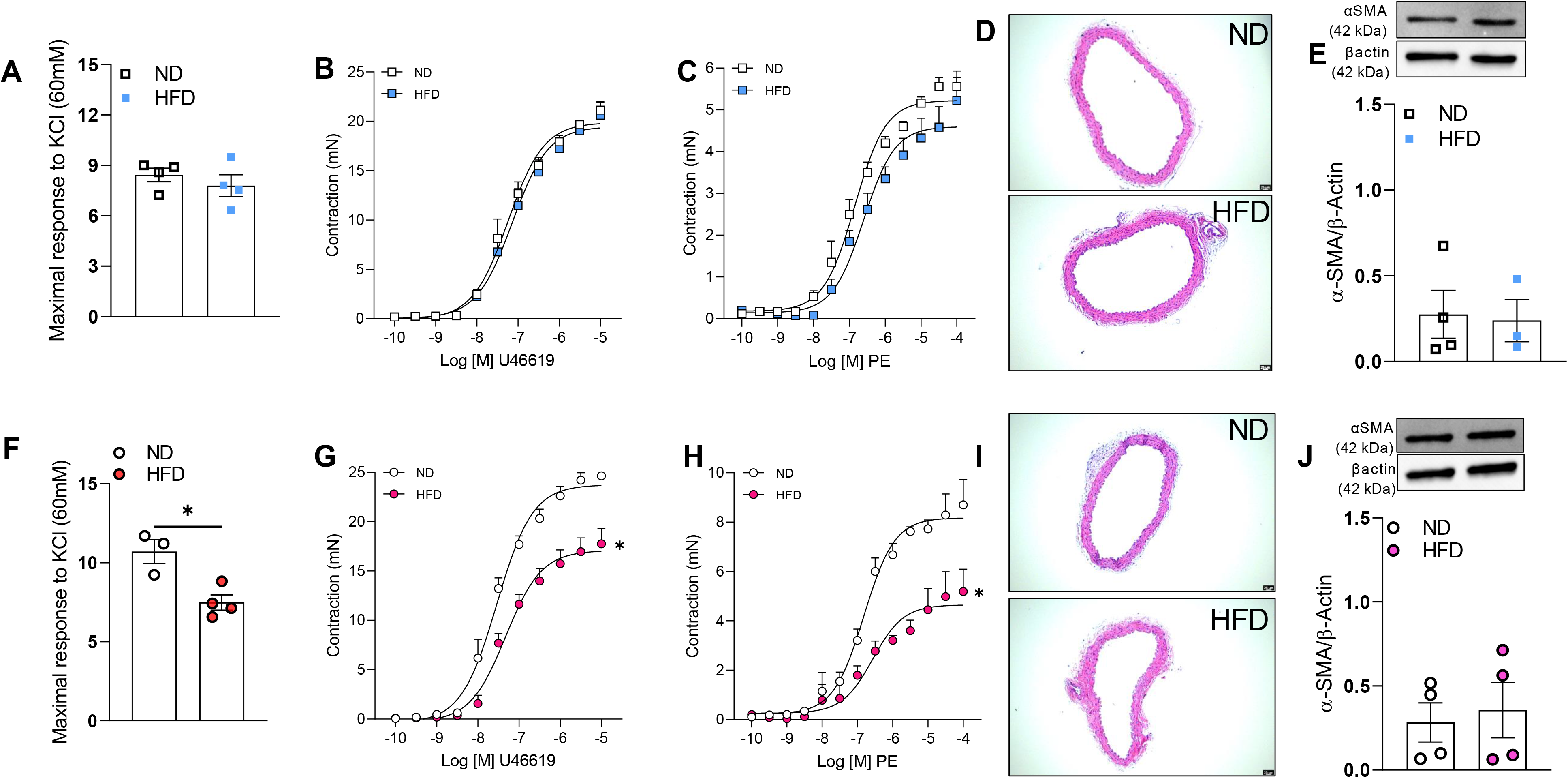
Vascular function and structure from male and female mice exposed to high fat diet (HFD). KCl, 60mMM response (A and F) and concentration response curve (CRC) to thromboxane analogue, U46619 (B and G) or phenylephrine (C and H) in endothelium intact aortic rings. Aortic remodeling (D and I) and aortic smooth muscle alpha-actin (αSMA) (E and J) expression. Experiments were performed in vascular samples from male and female mice exposed to normal diet (ND) or HFD for 12 weeks. Data are presented as mean ± Standard Error of the Mean (SEM). N=4. *P<0.05 vs normal diet (ND).

### Attenuated vascular contractility in obese female mice is mediated by a suppressed Rho-kinase activity

To study by which mechanism obese female mice display attenuated vascular contractility we inhibited Rho-kinase pathway via Y-27632. We observed that Y-27632 similarly affected the vascular contractility in arteries from lean and obese male mice (Fig. 6A) and Y-27632 triggered equal relaxation in lean and obese male mice (Fig. 6B). Furthermore, no difference in Mypt1 phosphorylation or total RhoA, ROCK1 and 2 was found in arteries from lean and obese male mice (Fig.6C-F).

**Figure 6.**
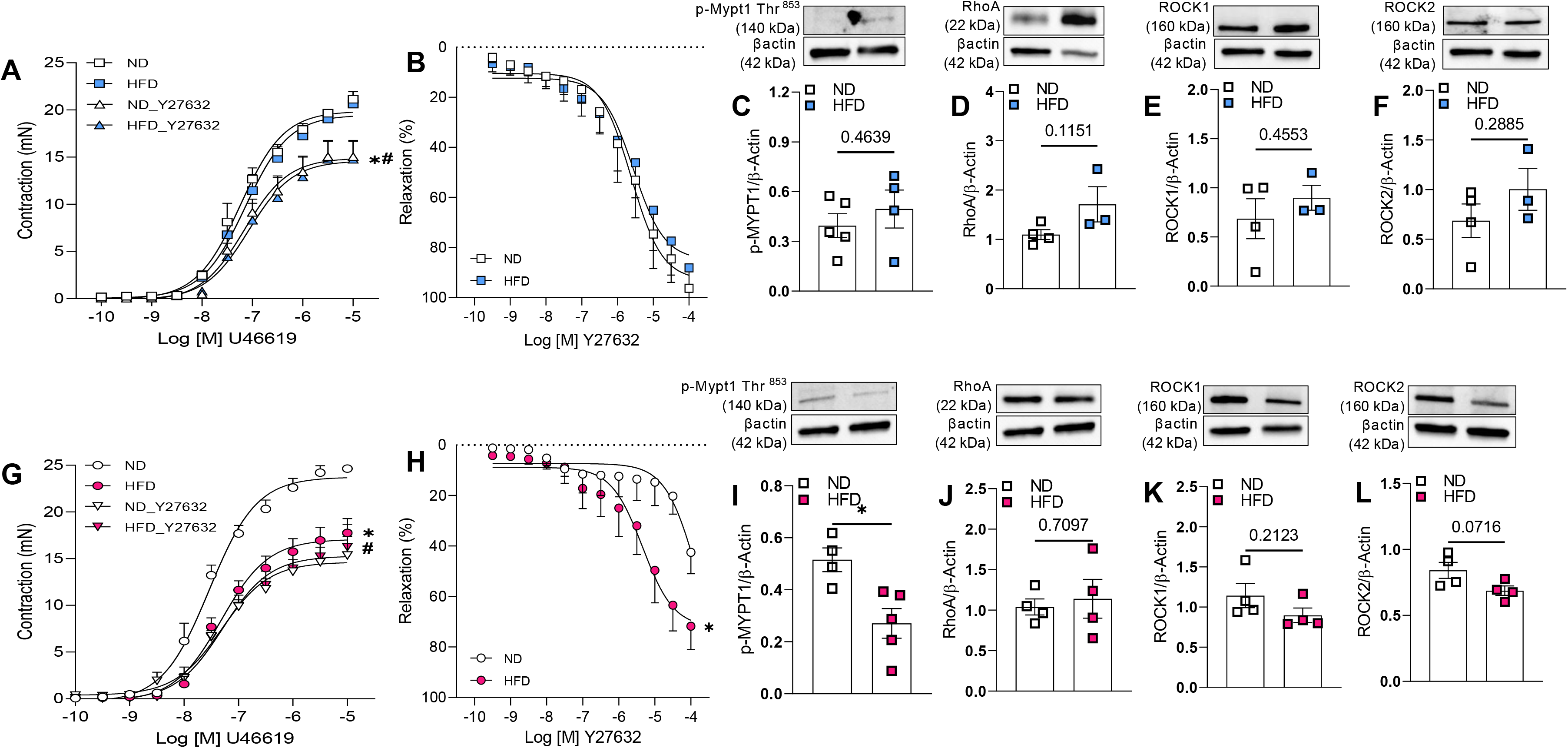
Role of Rho-kinase pathway on vascular dysfunction associated with high fat diet (HFD) treatment. Concentration response curve (CRC) to thromboxane analogue, U46619 with or without ROCK inhibitor (A and G) or CRC to Y27632 (B and H) in endothelium intact aortic rings. Expression of Rho-kinase pathway-associated proteins in aortae (C-F and I-L) analyzed by western blot. Experiments were performed in vascular samples from male and female mice exposed to normal diet (ND) or HFD for 12 weeks. Data are presented as mean ± Standard Error of the Mean (SEM). N=4. *P<0.05 vs normal diet (ND); ^#^P<0.05 vs without Y27632

We also observed that Y-27632 only affected the response of arteries from lean female mice, but not obese female mice (Fig.6G), suggesting that Rho-kinase pathway is attenuated in female mice exposed to HFD. To study the direct effect of Rho-kinase on modulating the vascular contractility in female mice, we built curves to Y-27632 and found increased maximal response in arteries from obese female mice (Fig. 6H). Finally, decreased phosphorylated Mypt1 at Thr^853^ residue, which is involved in RhoA/ROCK-mediated inhibition of myosin phosphatase^12^, was found in arteries from obese female mice (Fig. 6I), further confirming decreased Rho-kinase pathway. No difference was found for RhoA and ROCK1 and2 expression (Fig.6J-L).

## Discussion

In this study, we sought to describe the sex-specificity of the mechanisms controlling vascular contractility in obesity and pinpoint the source of any potential sex-discrepancy. Also, we looked into how body weight, metabolic issues, energy usage, and inflammation might be linked to vascular dysfunction. Our key findings are, (1) male and female mice develop characteristics of obesity HFD, but female mice are more resistant to HFD-induced body weight gain; (2) female mice present a better energy expenditure behavior under HFD; (3) aortae from obese female displayed hypocontractility, whereas male mice do not demonstrate any alteration; (4) finally female exposed to HFD show suppressed Rho-kinase activity. In light of these findings, we first established that female mice have lower Rho-kinase activation throughout the development of obesity as a potential compensatory strategy to safeguard the vasculature against obesity-related vascular damage.

HFD in rodents induces a sexual dimorphism in body weight, metabolic alterations, and degree of inflammation^22-26^. Female mice are commonly leaner and exhibit reduced increases in body weight, preserved metabolic function, and lower degree of inflammation as compared to male^24-27^. We previously demonstrated that male mice display a significant increase in body weight under obesogenic diet from week 3, whereas female only after week 9. In the same study, we observed that differences between male and female mice disappear only after 18 weeks of HFD intervention,^5^ indicating that female mice have a slower body weight gain under obesogenic diet, which might be associated with increased energy expenditure – since female demonstrated elevate heat in CLAMS analyze – therefore, an elevated energy burn could be a gatekeeper against the fat accumulation in female mice under obesogenic diet. Finally, we only treated our mice for 12-13 weeks with obesogenic diet, perhaps exposing the mice longer would blunt any sex difference at the end.

Fat accumulation can lead to impaired glucose response by promoting insulin resistance and disrupting glucose uptake and metabolism, which appears to be a sex-specific response. Male mice after 14 or 16 weeks of western diet^22^ or HFD^24, 27^ demonstrate a worse glucose metabolism compared to their female counterpart. On our hands, male and female mice became glucose tolerant, but male were more sensitive to obesogenic diet. Inflammation of adipose tissue is a key precursor of glucose tolerance^28, 29^, however female mice trend to present a lower inflammation then male under HFD^24, 30^. We observed that chemokine receptor CCR5, adhesion genes (VCAM1 and ICAM1), as well as F4/80 (macrophage marker) were upregulated in gonadal fat from obese male in at least 4-fold increases, in contrast female only displayed a mild increase in TNFα. CCR5 plays major role in controlling obesity-induced adipose tissue inflammation and insulin resistance by regulating macrophage recruitment^31^, therefore increased sensitivity to HFD-induced inflammation in male, likely dependent on CCR5 and macrophages, would justify why male become more glucose tolerant to HFD

We and others have demonstrated that obesity affects the function of large and small arteries^5, 16, 18, 32-34^ in a sex discrepancy-dependent manner^5, 7, 8, 22^. Although such information is already well-established, the molecular mechanisms is not fully comprehended. In this study we observed that only aortae from female exposed to obesogenic diet presented attenuated response to different contractile agonists, which was not associated with changes in vascular remodeling or contractile protein amount indicating that an intracellular signaling is altered only in female. Therefore, we investigated an important signaling pathway associated with cardiovascular risks^16, 35-38^, the Rho-kinase pathway.

Multiple vascular contractile agonists generate their responses by activating Rho-kinase pathway including endothelin-1^39^, angiotensin-II^40^, and arachidonic acid metabolites (thromboxane A_2_)^41^, in common, these important mediators are elevated in obesity^42-45^. We found that 1) inhibiting Rho-kinase signaling blunted the difference on vascular contraction between lean and obese female, 2) arteries from female mice exposed to obesogenic diet revealed a premature response to Y27632, and 3) Mypt-1 phosphorylation at Thr^853^ is attenuated in arteries from obese female mice, when taken together, these data imply that the Rho-kinase pathway is inhibited in the arteries of female mice given HFD. Many compensatory mechanisms in the vasculature of obese females have been proposed. Here, we propose for the first time that RhoA pathway blockage occurs in an effort to ensure a protection against obesity-induced cardiovascular risk in females. Previous findings suggest that estrogen suppresses Rho-kinase function in the cerebral circulation^46^, induces neuroprotective effects of in model of Parkinson’s disease via inhibiting Rho-kinase^47^, and causes a decrease in Rho-kinase mRNA expression^48^. Thus, we can suggest that increase in estrogen might be regulating the suppression of Rho-kinase pathway in obese female. Further investigations are necessary to confirm whether changes in estrogen signaling are driving the vascular protection in female mice and whether such adaptive response is endothelium dependent.

In summary, our data indicate that male mice are more susceptible to gain body weight compared to female mice, which is associated with impaired energy expenditure, higher glucose tolerance, and exacerbated adipose tissue inflammation. Our findings also suggest that female mice under obesogenic diet demonstrate a vascular protective effect by attenuating the Rho-kinase pathway, whereas male mice lack in any adaptive response.

Future studies will help to elucidate by which endocrine and vascular mechanisms female mice display such protection. Overall, our data add one more piece to the literature that sex should be considered an important variable when identifying the adequate therapeutic strategy for treatment of obesity associated vascular dysfunction as the therapies which are effective in one sex may not be effective in other

## Supporting information

Primers and antibodies

## Acknowledgement

This work was supported by The São Paulo Research Foundation (FAPESP, 2021/01069-0) to G.S.B and NHLBI-R00 (R00HL14013903) and startup funds from University of Pittsburgh to T.B.-N.

## References

1. Powell-Wiley TM, Poirier P, Burke LE, Despres JP, Gordon-Larsen P, Lavie CJ, Lear SA, Ndumele CE, Neeland IJ, Sanders P, St-Onge MP, American Heart Association Council on L, Cardiometabolic H, Council on C, Stroke N, Council on Clinical C, Council on e, Prevention and Stroke C. Obesity and Cardiovascular Disease: A Scientific Statement From the American Heart Association. Circulation. 2021;143:e984–e1010.

2. (WHO) WHO. Obesity and overweight. 2021;2023.

3. Berg AH and Scherer PE. Adipose tissue, inflammation, and cardiovascular disease. Circ Res. 2005;96:939–49.

4. Garawi F, Devries K, Thorogood N and Uauy R. Global differences between women and men in the prevalence of obesity: is there an association with gender inequality? Eur J Clin Nutr. 2014;68:1101–6.

5. Bruder-Nascimento T, Ekeledo OJ, Anderson R, Le HB and Belin de Chantemele EJ. Long Term High Fat Diet Treatment: An Appropriate Approach to Study the Sex-Specificity of the Autonomic and Cardiovascular Responses to Obesity in Mice. Front Physiol. 2017;8:32.

6. Faulkner JL, Bruder-Nascimento T and Belin de Chantemele EJ. The regulation of aldosterone secretion by leptin: implications in obesity-related cardiovascular disease. Curr Opin Nephrol Hypertens. 2018;27:63–69.

7. Victorio JA, Guizoni DM, Freitas IN, Araujo TR and Davel AP. Effects of High-Fat and High-Fat/High-Sucrose Diet-Induced Obesity on PVAT Modulation of Vascular Function in Male and Female Mice. Front Pharmacol. 2021;12:720224.

8. Davel AP, Lu Q, Moss ME, Rao S, Anwar IJ, DuPont JJ and Jaffe IZ. Sex-Specific Mechanisms of Resistance Vessel Endothelial Dysfunction Induced by Cardiometabolic Risk Factors. J Am Heart Assoc. 2018;7.

9. Faulkner JL, Kennard S, Huby AC, Antonova G, Lu Q, Jaffe IZ, Patel VS, Fulton DJR and Belin de Chantemele EJ. Progesterone Predisposes Females to Obesity-Associated Leptin-Mediated Endothelial Dysfunction via Upregulating Endothelial MR (Mineralocorticoid Receptor) Expression. Hypertension. 2019;74:678–686.

10. Link JC and Reue K. Genetic Basis for Sex Differences in Obesity and Lipid Metabolism. Annu Rev Nutr. 2017;37:225–245.

11. Reue K. Sex differences in obesity: X chromosome dosage as a risk factor for increased food intake, adiposity and co-morbidities. Physiol Behav. 2017;176:174–182.

12. Nunes KP, Rigsby CS and Webb RC. RhoA/Rho-kinase and vascular diseases: what is the link? Cell Mol Life Sci. 2010;67:3823–36.

13. Kolluru GK, Majumder S and Chatterjee S. Rho-kinase as a therapeutic target in vascular diseases: striking nitric oxide signaling. Nitric Oxide. 2014;43:45–54.

14. Takemoto K, Ishihara S, Mizutani T, Kawabata K and Haga H. Compressive stress induces dephosphorylation of the myosin regulatory light chain via RhoA phosphorylation by the adenylyl cyclase/protein kinase A signaling pathway. PLoS One. 2015;10:e0117937.

15. da Costa RM, Fais RS, Dechandt CRP, Louzada-Junior P, Alberici LC, Lobato NS and Tostes RC. Increased mitochondrial ROS generation mediates the loss of the anti-contractile effects of perivascular adipose tissue in high-fat diet obese mice. Br J Pharmacol. 2017;174:3527–3541.

16. Nguyen Dinh Cat A, Callera GE, Friederich-Persson M, Sanchez A, Dulak-Lis MG, Tsiropoulou S, Montezano AC, He Y, Briones AM, Jaisser F and Touyz RM. Vascular dysfunction in obese diabetic db/db mice involves the interplay between aldosterone/mineralocorticoid receptor and Rho kinase signaling. Sci Rep. 2018;8:2952.

17. Bruder-Nascimento T, Kennard S, Antonova G, Mintz JD, Bence KK and Belin de Chantemele EJ. Ptp1b deletion in pro-opiomelanocortin neurons increases energy expenditure and impairs endothelial function via TNF-alpha dependent mechanisms. Clin Sci (Lond). 2016;130:881–93.

18. da Costa RM, Neves KB, Mestriner FL, Louzada-Junior P, Bruder-Nascimento T and Tostes RC. TNFalpha induces vascular insulin resistance via positive modulation of PTEN and decreased Akt/eNOS/NO signaling in high fat diet-fed mice. Cardiovasc Diabetol. 2016;15:119.

19. Bruder-Nascimento T, Butler BR, Herren DJ, Brands MW, Bence KK and Belin de Chantemele EJ. Deletion of protein tyrosine phosphatase 1b in proopiomelanocortin neurons reduces neurogenic control of blood pressure and protects mice from leptin- and sympatho-mediated hypertension. Pharmacol Res. 2015;102:235–44.

20. Bruder-Nascimento T, Faulkner JL, Haigh S, Kennard S, Antonova G, Patel VS, Fulton DJR, Chen W and Belin de Chantemele EJ. Leptin Restores Endothelial Function via Endothelial PPARgamma-Nox1-Mediated Mechanisms in a Mouse Model of Congenital Generalized Lipodystrophy. Hypertension. 2019;74:1399–1408.

21. Bruder-Nascimento T, Kress TC, Pearson M, Chen W, Kennard S and Belin de Chantemele EJ. Reduced Endothelial Leptin Signaling Increases Vascular Adrenergic Reactivity in a Mouse Model of Congenital Generalized Lipodystrophy. Int J Mol Sci. 2021;22.

22. Padilla J, Woodford ML, Lastra-Gonzalez G, Martinez-Diaz V, Fujie S, Yang Y, Lising AMC, Ramirez-Perez FI, Aroor AR, Morales-Quinones M, Ghiarone T, Whaley-Connell A, Martinez-Lemus LA, Hill MA and Manrique-Acevedo C. Sexual Dimorphism in Obesity-Associated Endothelial ENaC Activity and Stiffening in Mice. Endocrinology. 2019;160:2918–2928.

23. Gupte M, Thatcher SE, Boustany-Kari CM, Shoemaker R, Yiannikouris F, Zhang X, Karounos M and Cassis LA. Angiotensin converting enzyme 2 contributes to sex differences in the development of obesity hypertension in C57BL/6 mice. Arterioscler Thromb Vasc Biol. 2012;32:1392–9.

24. Pettersson US, Walden TB, Carlsson PO, Jansson L and Phillipson M. Female mice are protected against high-fat diet induced metabolic syndrome and increase the regulatory T cell population in adipose tissue. PLoS One. 2012;7:e46057.

25. Ganz M, Csak T and Szabo G. High fat diet feeding results in gender specific steatohepatitis and inflammasome activation. World J Gastroenterol. 2014;20:8525–34.

26. Singer K, Maley N, Mergian T, DelProposto J, Cho KW, Zamarron BF, Martinez-Santibanez G, Geletka L, Muir L, Wachowiak P, Demirjian C and Lumeng CN. Differences in Hematopoietic Stem Cells Contribute to Sexually Dimorphic Inflammatory Responses to High Fat Diet-induced Obesity. J Biol Chem. 2015;290:13250–62.

27. Rudnicki M, Abdifarkosh G, Rezvan O, Nwadozi E, Roudier E and Haas TL. Female Mice Have Higher Angiogenesis in Perigonadal Adipose Tissue Than Males in Response to High-Fat Diet. Front Physiol. 2018;9:1452.

28. Burhans MS, Hagman DK, Kuzma JN, Schmidt KA and Kratz M. Contribution of Adipose Tissue Inflammation to the Development of Type 2 Diabetes Mellitus. Compr Physiol. 2018;9:1–58.

29. Wu H and Ballantyne CM. Metabolic Inflammation and Insulin Resistance in Obesity. Circ Res. 2020;126:1549–1564.

30. Chen KE, Lainez NM and Coss D. Sex Differences in Macrophage Responses to Obesity-Mediated Changes Determine Migratory and Inflammatory Traits. J Immunol. 2021;206:141–153.

31. Kitade H, Sawamoto K, Nagashimada M, Inoue H, Yamamoto Y, Sai Y, Takamura T, Yamamoto H, Miyamoto K, Ginsberg HN, Mukaida N, Kaneko S and Ota T. CCR5 plays a critical role in obesity-induced adipose tissue inflammation and insulin resistance by regulating both macrophage recruitment and M1/M2 status. Diabetes. 2012;61:1680–90.

32. Ferreira NS, Bruder-Nascimento T, Pereira CA, Zanotto CZ, Prado DS, Silva JF, Rassi DM, Foss-Freitas MC, Alves-Filho JC, Carlos D and Tostes RC. NLRP3 Inflammasome and Mineralocorticoid Receptors Are Associated with Vascular Dysfunction in Type 2 Diabetes Mellitus. Cells. 2019;8.

33. Bruder-Nascimento T, Callera GE, Montezano AC, He Y, Antunes TT, Nguyen Dinh Cat A, Tostes RC and Touyz RM. Vascular injury in diabetic db/db mice is ameliorated by atorvastatin: role of Rac1/2-sensitive Nox-dependent pathways. Clin Sci (Lond). 2015;128:411–23.

34. Rocha VDS, Claudio ERG, da Silva VL, Cordeiro JP, Domingos LF, da Cunha Mrh, Mauad H, do Nascimento TB, Lima-Leopoldo AP and Leopoldo AS. High-Fat Diet-Induced Obesity Model Does Not Promote Endothelial Dysfunction via Increasing Leptin/Akt/eNOS Signaling. Front Physiol. 2019;10:268.

35. Gabrielli L, Winter JL, Godoy I, McNab P, Padilla I, Cordova S, Rigotti P, Novoa U, Mora I, Garcia L, Ocaranza MP and Jalil JE. Increased rho-kinase activity in hypertensive patients with left ventricular hypertrophy. Am J Hypertens. 2014;27:838–45.

36. Masumoto A, Hirooka Y, Shimokawa H, Hironaga K, Setoguchi S and Takeshita A. Possible involvement of Rho-kinase in the pathogenesis of hypertension in humans. Hypertension. 2001;38:1307–10.

37. Denniss SG, Jeffery AJ and Rush JW. RhoA-Rho kinase signaling mediates endothelium- and endoperoxide-dependent contractile activities characteristic of hypertensive vascular dysfunction. Am J Physiol Heart Circ Physiol. 2010;298:H1391–405.

38. Bruder-Nascimento T, Callera G, Montezano AC, Antunes TT, He Y, Cat AN, Ferreira NS, Barreto PA, Olivon VC, Tostes RC and Touyz RM. Renoprotective Effects of Atorvastatin in Diabetic Mice: Downregulation of RhoA and Upregulation of Akt/GSK3. PLoS One. 2016;11:e0162731.

39. Lima VV, Giachini FR, Carneiro FS, Carvalho MH, Fortes ZB, Webb RC and Tostes RC. O-GlcNAcylation contributes to the vascular effects of ET-1 via activation of the RhoA/Rho-kinase pathway. Cardiovasc Res. 2011;89:614–22.

40. Kimura K and Eguchi S. Angiotensin II type-1 receptor regulates RhoA and Rho-kinase/ROCK activation via multiple mechanisms. Focus on “Angiotensin II induces RhoA activation through SHP2-dependent dephosphorylation of the RhoGAP p190A in vascular smooth muscle cells”. Am J Physiol Cell Physiol. 2009;297:C1059–61.

41. Wilson DP, Susnjar M, Kiss E, Sutherland C and Walsh MP. Thromboxane A2-induced contraction of rat caudal arterial smooth muscle involves activation of Ca2+ entry and Ca2+ sensitization: Rho-associated kinase-mediated phosphorylation of MYPT1 at Thr-855, but not Thr-697. Biochem J. 2005;389:763–74.

42. Weil BR, Westby CM, Van Guilder GP, Greiner JJ, Stauffer BL and DeSouza CA. Enhanced endothelin-1 system activity with overweight and obesity. Am J Physiol Heart Circ Physiol. 2011;301:H689–95.

43. White MC, Miller AJ, Loloi J, Bingaman SS, Shen B, Wang M, Silberman Y, Lindsey SH and Arnold AC. Sex differences in metabolic effects of angiotensin-(1-7) treatment in obese mice. Biol Sex Differ. 2019;10:36.

44. Saiki A, Ohira M, Endo K, Koide N, Oyama T, Murano T, Watanabe H, Miyashita Y and Shirai K. Circulating angiotensin II is associated with body fat accumulation and insulin resistance in obese subjects with type 2 diabetes mellitus. Metabolism. 2009;58:708–13.

45. Graziani F, Biasucci LM, Cialdella P, Liuzzo G, Giubilato S, Della Bona R, Pulcinelli FM, Iaconelli A, Mingrone G and Crea F. Thromboxane production in morbidly obese subjects. Am J Cardiol. 2011;107:1656–61.

46. Chrissobolis S, Budzyn K, Marley PD and Sobey CG. Evidence that estrogen suppresses rho-kinase function in the cerebral circulation in vivo. Stroke. 2004;35:2200–5.

47. Rodriguez-Perez AI, Dominguez-Meijide A, Lanciego JL, Guerra MJ and Labandeira-Garcia JL. Inhibition of Rho kinase mediates the neuroprotective effects of estrogen in the MPTP model of Parkinson’s disease. Neurobiol Dis. 2013;58:209–19.

48. Hiroki J, Shimokawa H, Mukai Y, Ichiki T and Takeshita A. Divergent effects of estrogen and nicotine on Rho-kinase expression in human coronary vascular smooth muscle cells. Biochem Biophys Res Commun. 2005;326:154–9.

