## Supplementary material for "Suppressed vascular Rho-kinase activation is a protective cardiovascular mechanism in obese female mice": Primers and antibodies

**Supplemental Table 1. List of primers**

| **Primer** | **Sequence** | |
|---|---|---|
| CCR1 | FW | GCCAAAAGACTGCTGTAAGAGCC |
| | RV | GCTTTGAAGCCTCCTATGCTGC |
| CCR3 | FW | CCACTGTACTCCCTGGTGTTCA |
| | RV | GGACAGTGAAGAGAAAGAGCAGG |
| CCR5 | FW | GGTTCCTGAAAGCGGCTGTAAATA |
| | RV | CTGTTGGCAGTCAGGCACATC |
| F4/80 | FW | TCCTGCTGTGTCGTGCTGTTC |
| | RV | GCCGTCTGGTTGTCAGTCTTGTC |
| IL6 | FW | TTCTTGGGACTGCTGGT |
| | RV | CAGGTCTGTTGGGAGTGGTA |
| TNF[illegible] | FW | AATGGCCTCCCTCTCATCAG |
| | RV | CCTAACTGCCCTTCCTCCAT |
| Ki67 | FW | AGAGCCTTAGCAATAGCAACG |
| | RV | GTCTCCCGCGATTCCTCTG |
| VCAM1 | FW | TGACAAGTCCCCATCGTTGA |
| | RV | ACCTCGCGACGGCATAATT |
| ICAM1 | FW | ATCACATGGGTCGAGGGTTT |
| | RV | AACCACTGCCAGTCCACATA |
| GAPDH | FW | GAGAGGCCCTATCCCAACTC |
| | RV | TCAAGAGAGTAGGGAGGGCT |

Primers were purchased from Integrated DNA Technologies

**Supplemental Table 2. List of antibodies**

| Antibody | Catalog number | Company | Concentration |
|---|---|---|---|
| αSMA | 19245 | Cell Signaling | 1:2000 |
| Thr 853 Mypt1 | 4563 | Cell Signaling | 1:500 |
| RhoA | 2117 | Cell Signaling | 1:1000 |
| ROCK1 | 28999 | Cell Signaling | 1:1000 |
| ROCK2 | 47012 | Cell Signaling | 1:1000 |
| βactin | A3854 | Sigma | 1:20000 |
